# Fast, sensitive, and accurate integration of single cell data with Harmony

**DOI:** 10.1101/461954

**Authors:** Ilya Korsunsky, Jean Fan, Kamil Slowikowski, Fan Zhang, Kevin Wei, Yuriy Baglaenko, Michael Brenner, Po-Ru Loh, Soumya Raychaudhuri

## Abstract

The rapidly emerging diversity of single cell RNAseq datasets allows us to characterize the transcriptional behavior of cell types across a wide variety of biological and clinical conditions. With this comprehensive breadth comes a major analytical challenge. The same cell type across tissues, from different donors, or in different disease states, may appear to express different genes. A joint analysis of multiple datasets requires the integration of cells across diverse conditions. This is particularly challenging when datasets are assayed with different technologies in which real biological differences are interspersed with technical differences. We present Harmony, an algorithm that projects cells into a shared embedding in which cells group by cell type rather than dataset-specific conditions. Unlike available single-cell integration methods, Harmony can simultaneously account for multiple experimental and biological factors. We develop objective metrics to evaluate the quality of data integration. In four separate analyses, we demonstrate the superior performance of Harmony to four single-cell-specific integration algorithms. Moreover, we show that Harmony requires dramatically fewer computational resources. It is the only available algorithm that makes the integration of *∼* 10^6^ cells feasible on a personal computer. We demonstrate that Harmony identifies both broad populations and fine-grained subpopulations of PBMCs from datasets with large experimental differences. In a meta-analysis of 14,746 cells from 5 studies of human pancreatic islet cells, Harmony accounts for variation among technologies and donors to successfully align several rare subpopulations. In the resulting integrated embedding, we identify a previously unidentified population of potentially dysfunctional alpha islet cells, enriched for genes active in the Endoplasmic Reticulum (ER) stress response. The abundance of these alpha cells correlates across donors with the proportion of dysfunctional beta cells also enriched in ER stress response genes. Harmony is a fast and flexible general purpose integration algorithm that enables the identification of shared fine-grained subpopulations across a variety of experimental and biological conditions.

---

Recent technological advances^1^ have enabled unbiased single cell transcriptional profiling of thousands of cells in a single experiment. Projects such as the Human Cell Atlas^2^ (HCA) and Accelerating Medicines Partnership^3, 4^ exemplify the growing body of reference datasets of primary human tissues. While individual experiments contribute incrementally to our understanding of cell types, a comprehensive catalogue of healthy and diseased cells will require the integration of multiple datasets across donors, studies, and technological platforms. Moreover, in translational research, joint analyses across tissues and clinical conditions will be essential to identify disease expanded populations. However, meaningful biological variation in single cell RNA-seq datasets from different studies is often hopelessly confounded by data source.^5^ Recognizing this key issue, investigators have developed unsupervised multi-dataset integration algorithms, such as Seurat MultiCCA,^6^ MNN Correct,^7^ Scanorama,^8^ and BBKNN^9^ to enable joint analysis. These methods embed cells from diverse experimental conditions and biological contexts into a common reduced dimensional embedding to enable shared cell type identification across datasets.

Here we introduce Harmony, an algorithm for robust, scalable, and flexible multi-dataset integration that addresses, to meet three key challenges of unsupervised scRNAseq joint embedding. First, cell types with regulatory or pathogenic roles are often rare, with subtle transcriptomic signatures. Integration must be able to identify both common and rare cell types, particularly those whose subtle signatures are initially obscured by technical or biological confounders. To be sensitive to subpopulations with subtle signatures, Harmony uses a two-step iterative strategy that removes the effect of such confounding factors at each round. This makes it easier to identify shared cell types whose expression signatures were obscured in the original data. Second, the number of cells in experiments is quickly expanding, exceeding 100,000 cells in atlas-like datasets. Integration algorithms must scale computationally, both in terms of runtime and required memory resources. To scale to big data, Harmony uses linear methods and avoids the costly cell-to-cell comparisons that scale quadratically with the number of cells. Third, more complex experimental design of single cell analysis compares cells from different donors, tissues, and technological platforms. In order for the joint embedding to be free of the influence of each, integration must simultaneously account for multiple sources of variation. The Harmony simultaneously accounts for multiple sources of variation, because the clustering objective function is formulated to account for any number of categorical covariates. Harmony is available as an R package on github (https://github.com/immunogenomics/harmony), with functions for standalone and Seurat6 pipeline analyses.

Here, we demonstrate how Harmony address the three unmet needs outlined above. First, we give an overview of the Harmony algorithm. Then, we integrate carefully designed cell line datasets to introduce metrics for quantifying cell-type accuracy and degree of dataset-mixing before and after integration. All measures of cell-type accuracy are based on annotation within datasets separately. As Harmony does not use cell-type information to integrate cells, these labels provide an unbiased quantification of accuracy. We then demonstrate that Harmony generates embeddings with higher quality and fewer computational resources than all currently available algorithms by testing each of them across a wide range of dataset sizes, from 25,000 to 500,000 total cells. Next, we show that Harmony to identify both broad populations and fine-grained sub-populations of cells in three datasets of peripheral blood mononuclear cell (PBMC) datasets with large technical differences. Finally, in a pancreatic islet cell meta-analysis, we demonstrate the power of Harmony to simultaneously integrate donor-and technology-specific effects to identify several rare subpopulations, including one putative novel islet cell subtype.

## Results

### Harmony Iteratively Learns a Cell-Specific Linear Correction Function

Starting with a PCA embedding, Harmony first groups cells into multi-dataset clusters (**Figure 1A**). We use soft clustering, assigning cells to potentially multiple clusters, to account for smooth transitions between cell states. In our thinking, these clusters serve as surrogate variables, rather than actually defining discrete cell-types. We developed a novel variant of soft k-means clustering to favor clusters with representation across multiple datasets (Online Methods). Clusters containing cells that are disproportionately represented by a small subset of datasets are penalized by an information theoretic metric. Harmony allows the user to apply multiple different penalties to accommodate multiple technical or biological factors, such as different batches and different technology platforms. The uncertainty encoded in the soft clustering preserves discrete and continuous topologies while avoiding local minima that might result from too quickly maximizing representation across multiple datasets, and preserves uncertainty. After clustering, each dataset has a cluster-specific centroid (**Figure 1B**) that is used to compute cluster-specific linear correction factors (**Figure 1C**). Under favorable conditions, the surrogate variables, defined by cluster membership, correspond to cell types and cell states. Thus, the cluster-specific correction factors that Harmony computes correspond to individual cell-type and cell-state specific correction factors. In this way, Harmony learns a simple linear adjustment function that is sensitive to intrinsic cellular phenotypes. Finally, each cell is assigned a cluster-weighted average of these terms and corrected by its cell-specific linear factor (**Figure 1D**). As a result, each cell has a potentially unique correction factor, depending on its soft clustering distribution. Harmony iterates these four steps until convergence. At convergence, additional iterations would assign cells to the same clusters and compute the same linear correction factors.

**Figure 1.**
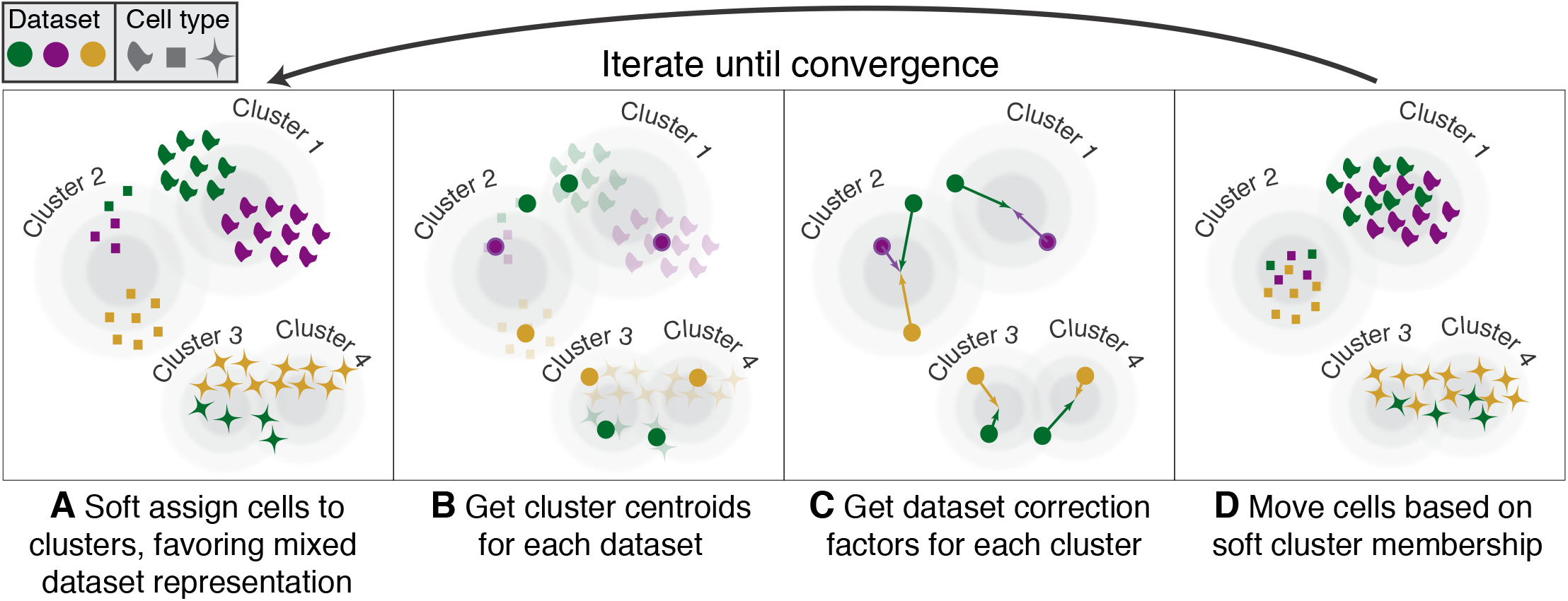
Overview of Harmony algorithm. We represent datasets with colors, and different cell types with shapes. Before we apply Harmony, principal components analysis embeds cells into a space with reduced dimensionality. Harmony accepts the cell coordinates in this reduced space and runs an iterative algorithm to adjust for data set specific effects. (A) Harmony uses fuzzy clustering to assign each cell to multiple clusters, while a penalty term ensures that the diversity of datasets within each cluster is maximized. (B) Harmony calculates a global centroid for each cluster, as well as dataset-specific centroids for each cluster. (C) Within each cluster, Harmony calculates a correction factor for each dataset based on the centroids. (D) Finally, Harmony corrects each cell with a cell-specific factor: a linear combination of dataset correction factors weighted by its soft cluster assignments made in step A. Harmony repeats steps A through D until convergence. The dependence between cluster assignment and dataset diminishes with each round.

### Quantification of Error and Accuracy in Cell Line Integration

We first assessed Harmony using three carefully controlled datasets, in order to evaluate performance on both integration (mixing of datasets) and accuracy (no mixing of cell types). Both metrics are important. Perfect integration can be achieved by simply mixing all cells, regardless of cellular identity. Similarly, high accuracy can be achieved by partitioning cell types into broad clusters without mixing datasets in small neighborhoods. In this situation, broad cellular states are defined, but fine-grained cellular substates and subtypes are confounded by the originating dataset. In order to quantify integration and accuracy of this embedding we felt that it was important that we have an objective metric. To this end, we compute the Local Inverse Simpson’s Index (LISI, Online Methods) in the local neighborhood of each cell. To assess integration, we employ integration LISI (iLISI, **Figure 2A**), denotes the effective number of datasets in a neighborhood. Neighborhoods represented by only a single dataset get an iLISI of 1, while neighborhoods with an equal number of cells from 2 datasets get an iLISI of 2. Note that even under ideal mixing, if the datasets have different numbers of cells, iLISI would be less than 2. To assess accuracy, we use cell-type LISI (cLISI, **Figure 2B**), the same mathematical measure, but applied to cell-type instead of dataset labels. Accurate integration should maintain a cLISI of 1, reflecting a separation of unique cell types throughout the embedding. An erroneous embedding would include neighborhoods with a cLISI of 2, indicating that neighbors have 2 different types of cells.

**Figure 2:**
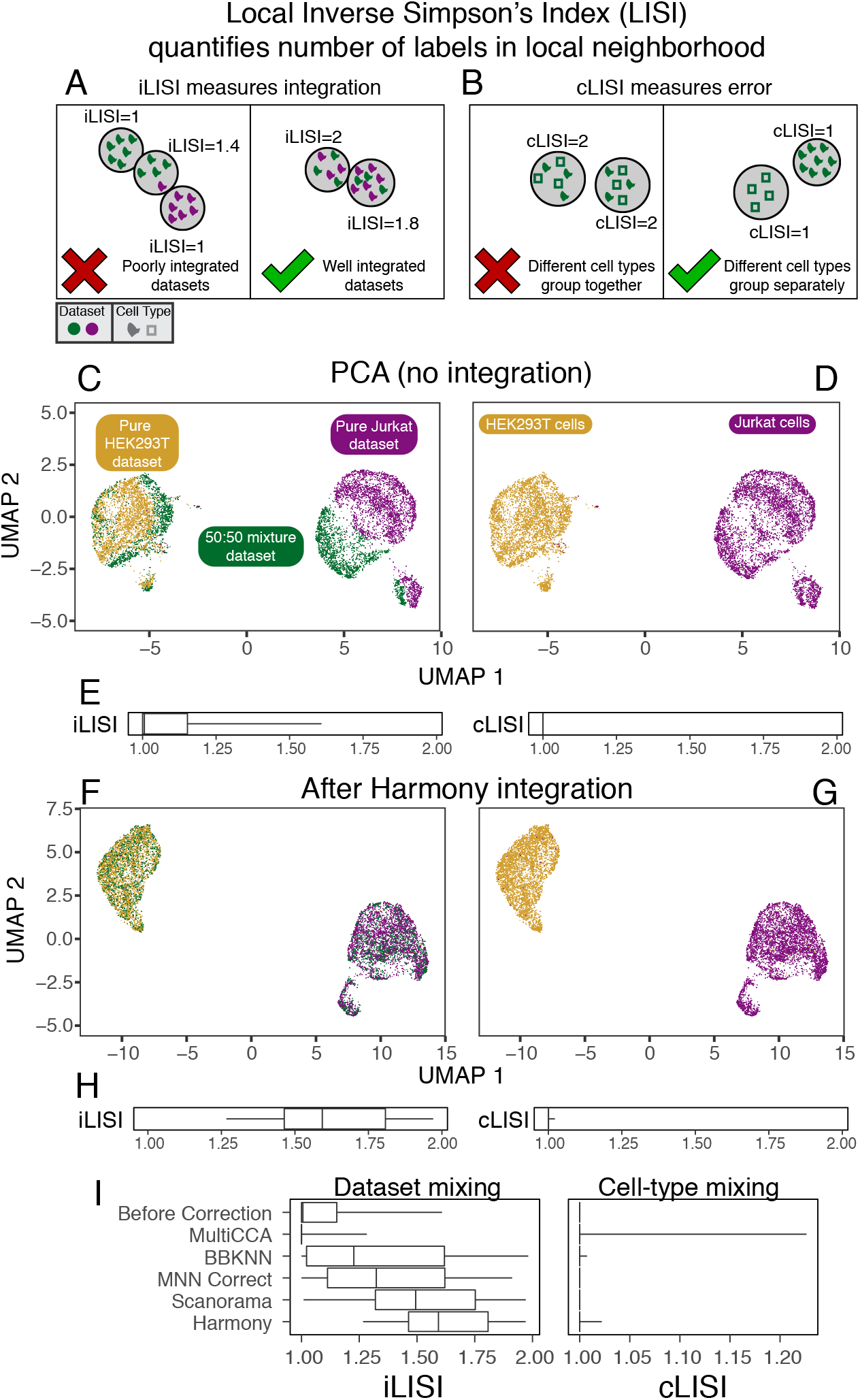
Quantitative assessment of dataset mixing and cell-type accuracy with cell line datasets. (A) iLISI measures the degree of mixing among datasets in an embedding, ranging from 1 in an unmixed space to B in a well mixed space. B is the number of datasets in the analysis. (B) cLISI measures integration accuracy using the same formulation but computed on cell-type labels instead. An accurate embedding has a cLISI close to 1 for every neighborhood, reflecting separation of different cell types. Jurkat and HEK293T cells from pure (purple and yellow) and mixed (green) cell-line datasets were analyzed together. Before Harmony integration, cells grouped by dataset (C) and known cell-type (D). iLISI and cLISI (E) were computed for every cell’s neighborhood and summarized with quantiles (5, 25, 50, 75, 95). After Harmony integration, cells from the mixture dataset are mixed into the other datasets (F), achieved by mixing Jurkat with Jurkat cells and HEK293T with HEK293T cells (G). iLISI and cLISI (H) were re-computed in the Harmony embedding. (I) This analysis was repeated for other algorithms and compared against no integration and Harmony integration using iLISI and cLISI quantiles.

We begin with three datasets from two cells lines: (1) pure Jurkat, (2) pure 293T and (3) a 50:50 mix.^10^ These datasets are particularly ideal for illustration and for assessment, as each cell can be unambiguously labeled Jurkat or 293T (**Figure S1**). A thorough integration would mix the 1799 Jurkat cells from the mixture dataset with 3255 cells from the pure Jurkat dataset and the 1565 293T cells from the mixture dataset with the 2859 from the pure 293T dataset. Thus, we expect the average iLISI to range from 1, reflecting no integration, to 1.8 for Jurkat cells and 1.5 for 293T cells^1^, reflecting maximal accurate integration. Application of a standard PCA pipeline followed by UMAP embedding demonstrates that the cells group broadly by dataset and cell type. This is both visually apparent (**Figure 2C,D**) and quantified (**Figure 2E,F**) with high accuracy reflected by a low cLISI (median iLISI 1.00, 95% [1.00, 1.00]). However the iLISI (median iLISI 1.01, 95% [1.00, 1.61]) is also low, reflecting imperfect integration, and ample structure within each cell-type reflecting the data set of origin. After Harmony, cells from the 50:50 dataset are mixed into the pure datasets (**Figure 2E**), which is appropriate in this case since these cell-lines have no additional biological structure. The increased iLISI (**Figure 2D**, median iLISI 1.59, 95% [1.27, 1.97]) reflects the mixing of datasets, while the low cLISI (median iLISI 1.00, 95% [1.00, 1.02]) reflects the accurate separation of Jurkat from 293T cells. iLISI and cLISI provide a quantitative way to assess the integration and accuracy of multiple algorithms. We repeated the integration and LISI analyses with MNN Correct, BBKNN, MultiCCA, and Scanorama (**Figure 2G**). While Harmony had the highest iLISI, Scanorama MNN Correct, and BBKNN provided lower levels of integration, evidence by lower iLISI. MultiCCA actually separated previously mixed datasets, yielding a lower iLISI (median 1.00, 95% [1.00, 1.28]) than before integration. Except for MultiCCA (median cLISI 1.00, 95% [1.00, 1.23]), the other algorithms maintained high accuracy (**Table S1**, median cLISI 1.00, 95% [1.00, 1.00]). This benchmark demonstrates the two key metrics for assessing mixing and accuracy and shows that Harmony performs well on both metrics in a well-controlled analysis of cell-line datasets.

### Harmony Scales to Enable Analysis of Large Data

As an integral part of the scRNAseq analysis pipeline, the integration algorithm must run in a reasonable amount of time and within the memory constraints of standard computers. To this end, we evaluated the computational performance of Harmony, measuring both the total runtime and maximum memory usage. To demonstrate the scalability of Harmony versus other methods, we downsampled from HCA data^11^ (528,688 cells from 16 donors and 2 tissues) to create 5 benchmark datasets with 500,000, 250,000, 125,000, 60,000, and 30,000 cells. We reported the runtime (**Table S2**) and memory (**Table S3**) for all benchmarks. Harmony runtime scaled well for all datasets (**Figure 3A**), ranging from 4 minutes on 30,000 cells to 68 minutes on 500,000 cells, 30 to 200 times faster than MultiCCA and MNN Correct. The runtimes for Harmony, BBKNN, and Scanorama were comparable for datasets with up to 125,000 cells. Harmony required very little memory (**Figure 3B**) compared to other algorithms, only 0.9GB on 30,000 cells and 7.2GB on 500,000 cells. At 125,000 cells, Harmony required 30 to 50 times less memory than Scanorama, MNN Correct and Seurat MultiCCA; these other methods could not scale beyond 125,000 cells. Notably, BBKNN was the only other algorithm able to finish on the 500,000 cell dataset, taking 44 minutes and 45GB of RAM. However, the BBKNN embedding did not achieve better integrated of tissues (**Figure 3C, S2A**, median iLISI 1.00, 95% [1.00, 1.10]) or donors (**Figure 3D, S2B**, median iLISI 1.60, 95% [1.00, 4.11]) above PCA alone (**Figure S2C** tissue median iLISI 1.00, 95% [1.00, 1.03], **Figure S2D** donor median iLISI 1.38, 95% [1.00, 3.14]).

**Figure 3:**
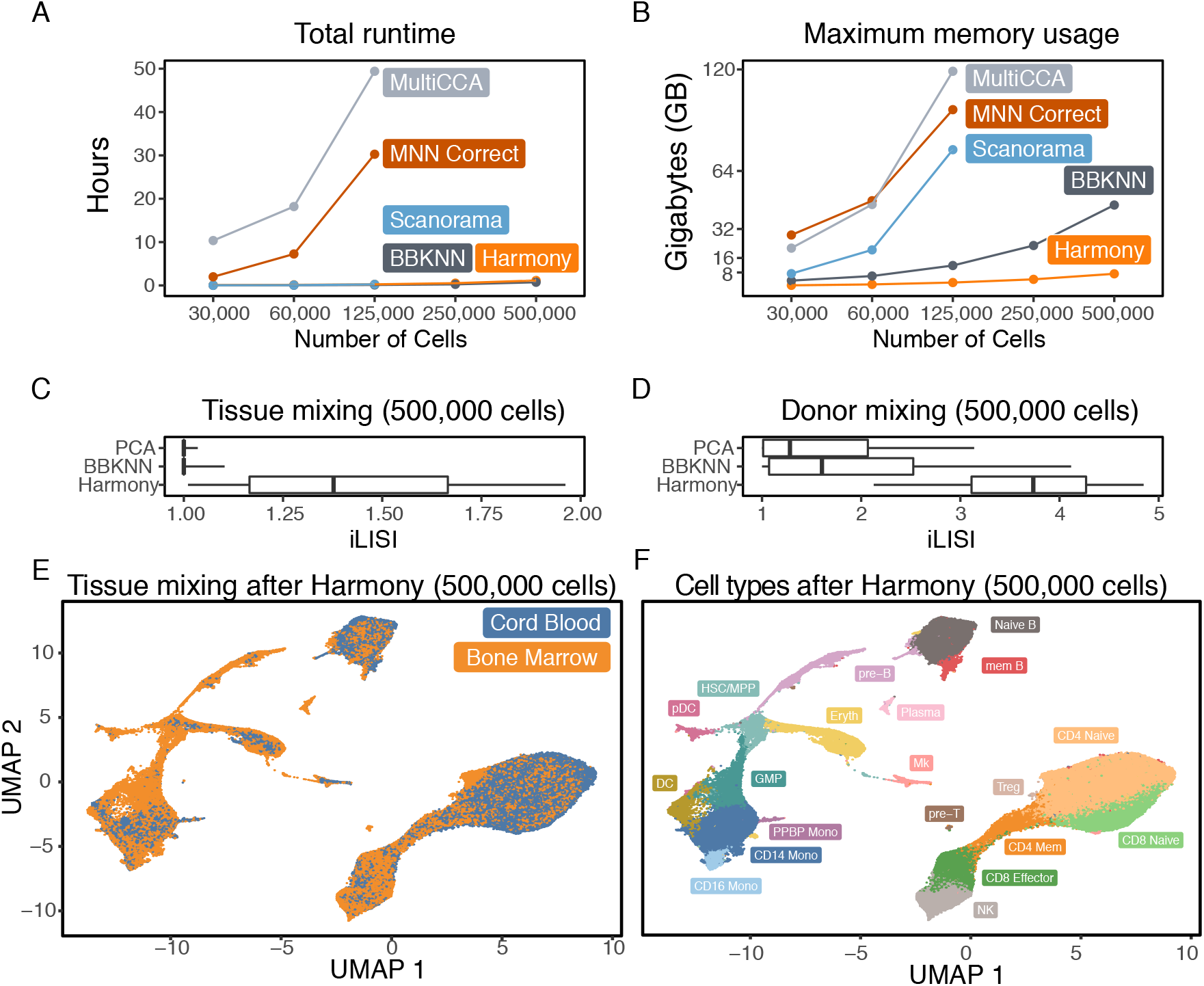
Computational efficiency benchmarks. We ran Harmony, BBKNN, Scanorama, MNN Correct, and MultiCCA on 5 downsampled HCA datasets of increasing sizes, from 25,000 to 500,000 cells. We recorded the (A) total runtime and (B) maximum memory required to analyze each dataset. Scanorama, MultiCCA, and MNN Correct were terminated for excessive memory requests on the 250,000 and 500,000 cell datasets. For Scanorama and Harmony, we quantified the extent of integration in the 500,000 cell benchmark for (C) the two tissues (cord blood and bone marrow) and (D) the 16 donors. For reference, (C) and (D) report the iLISI scores prior to integration. The mixing between tissues in the Harmony embedding is visualized in (E). In the Harmony embedding, (F) we clustered cells and labeled populations by canonical markers: pre-T cells, CD4 Naive T cells, CD4 Memory T cells, T-regs, CD8 Naive T cells, CD8 Effector T cells, natural killer cells (NK), pre-B cells, Naive B cells, Memory B cells, plasma cells, plasmacytoid dendritic cells (pDC), conventional dendritic cells (DC), granulocyte macrophage progenitor (GMP), CD16-monocytes (CD14 Mono), CD16+ monocytes (CD16 Mono), a population of monocytes also positive for Megakaryocyte markers (PPBP Mono), Megakaryocytes (Mk), Erythroid progenitors (Eryth), and a cluster of hematopoietic stem cells and multipotent progenitor cells (HSC/MPP).

Importantly, in addition to better computational performance of other algorithms Harmony returned a substantially more integrated space that other competing algorithms, allowing for the identification of shared cell types across tissues (**Figure 3E**, **Table S4**, median iLISI 1.40, 95% [1.04, 1.97] compared to medians of 1.00 to 1.12) and donors (**Figure S3**, median iLISI 3.93, 95% [2.46, 4.95] compared to medians of 1.07 to 2.82). In the Harmony embedding, we clustered cells and were able to identify shared populations (**Figure 3F**) using canonical markers (**Figure S4**). These results demonstrate that Harmony is computationally efficient and capable of analyzing even large datasets (10^5^ – 10^6^ cells) on personal computers. Alternative methods may require extensive parallelization to run modestly sized datasets.

### Deep Integration Enables Identification of Broad and Fine-Grained PBMCs Subpopulations

To assess how effective Harmony might be under more challenging scenarios, we gathered three datasets of human PBMCs, each assayed on the Chromium 10X platform but prepared with different protocols: 3-prime end v1 (3pV1), 3-prime end v2 (3pV2), and 5-prime (5p) end chemistries. After pooling all the cells together, we performed a joint analysis. Before integration, cells group primarily by dataset (**Figure 4A**, median iLISI 1.00, 95% [1.00, 1.00]). Harmony integrates the three datasets (**Figure 4B**, median iLISI 1.96, 95% [1.36, 2.56]), more than other methods (**Figure 4C**). To assess accuracy, within each dataset, we separately annotated (online methods) broad cell clusters with canonical markers of major expected populations (**Figure S5**): monocytes (*CD14*+ or *CD16*+), dendritic cells (*FCER1A*+), B cells (*CD20*+), T cells (*CD3*+), Megakaryocytes (*PPBP*+), and NK cells (*CD3*-/*GNLY*+) before clustering. We observed that Harmony retained differences among cell types (**Figure 4D** median cLISI 1.00, 95% [1.00, 1.02]). The greater dataset integration, compared to other algorithms, affords a unique opportunity to identify fine-grained cell subtypes. Using canonical markers (**Figure 4E**), we identified shared subpopulations of cells (**Figure 4F**) including naive CD4 T (*CD4*+/*CCR7*+), effector memory CD4 T (*CD4*+/*CCR7*-), Treg (*CD4*+/*FOXP3*+), memory CD8 (*CD8*+/*GZMK*-), effector CD8 T (*CD8*+/*GZMK*+), naive B (*CD20*+/*CD27*-), and memory B cells (*CD20*+/*CD27*+). In the embeddings produced by other algorithms, the median iLISI did not exceed 1.1 (**Table S5**). Accordingly, the subtypes identified with Harmony reside in dataset-specific, rather than dataset-mixed clusters (**Figure S6**). These results show that Harmony successfully accounts for technical variation among different protocols and integrates many different cell types while preserving large-scale and fine-grained structures in the data.

**Figure 4:**
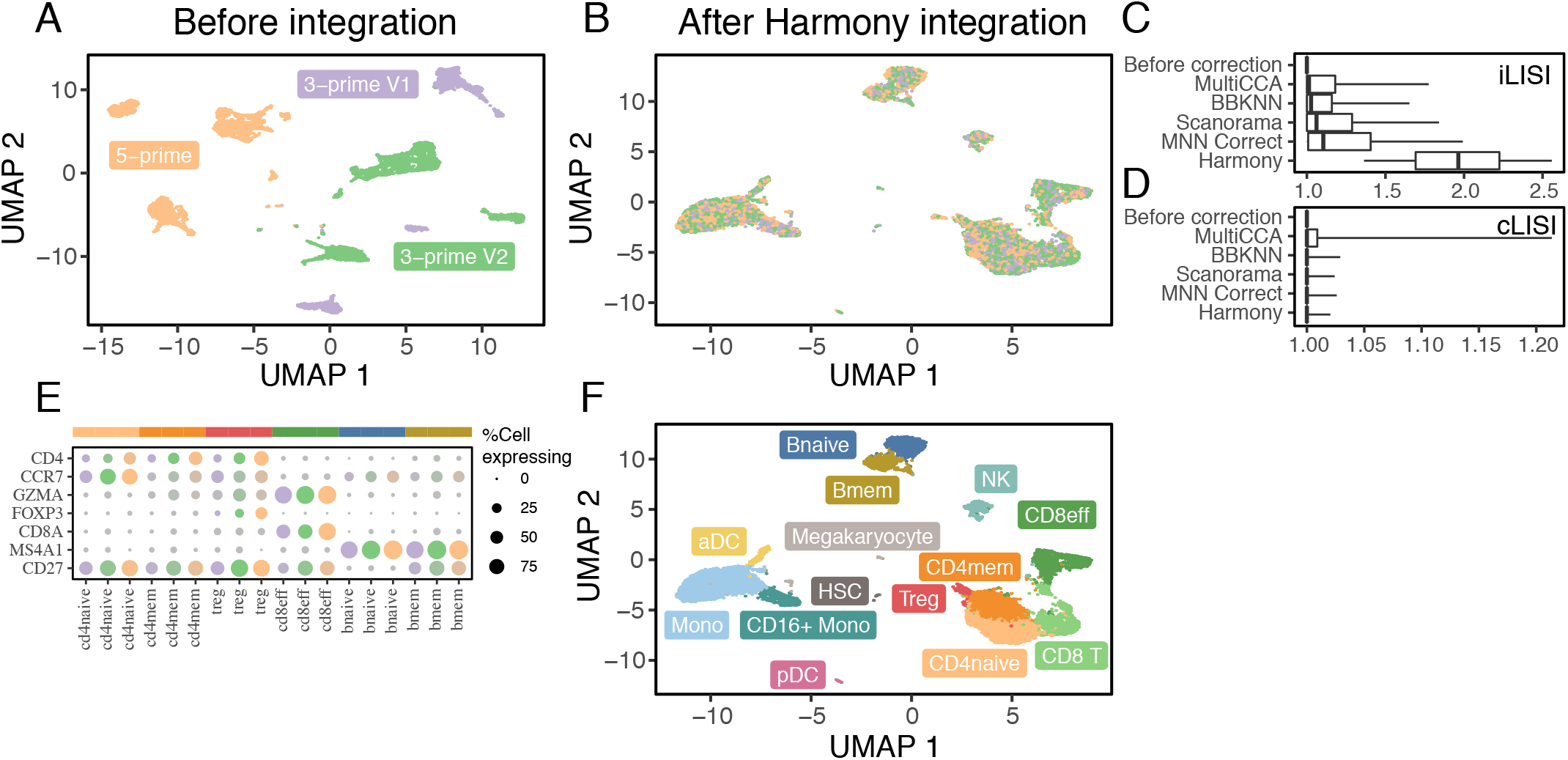
Fine-grained subpopulation identification in PBMCs across technologies. Three PBMC datasets were assayed with 10X, using different library construction protocols: 5-prime (orange), 3-prime V1 (purple), and 3-prime V2 (green). Before integration (A), cells group by dataset. After Harmony integration (B), datasets are mixed together. (C) Harmony achieves the most thorough integration among datasets, while preserving (D) cell type differences. Using canonical markers (E), we identified (F) 5 shared subtypes of T cells and 2 shared subtypes of B cells. (G) Other integration algorithms fail to group these cells by subtype.

### Simultaneous Integration Across Donors and Technologies Identifies Rare Pancreas Islet Subtypes

We considered a more complex experimental design, in which integration must be performed simultaneously over more than one variable. We gathered human pancreatic islet cells from independent five studies^12–16^, each of which were generated with a different technological platform. Integration across platforms is already challenging. However, within two^12, 13^ of the datasets, the authors also reported significant donor-specific effects. In this scenario, a successful integration of these studies must account for the effects of both technologies and donors, which may both affect different cell types in different ways. Harmony is the only single cell integration algorithm that is able to explicitly integrate over more than one variable, hence we omit a comparison against other methods.

As before, we assess cell type accuracy cLISI with canonical cell types identified independently within each dataset (**Figure S7**): alpha (*GCG*+), beta (*MAFA*+), gamma (*PPY*+), delta (*SST*+), acinar (*PRSS1*+), ductal (*KRT19*+), endothelial (*CDH5*+), stellate (*COL1A2*+), and immune (*PTPRC*+). Since there are two integration variables, we asses both donor iLISI and technology iLISI. Prior to integration, PCA separates cells by technology (**Figure 5A,E**, median iLISI 1.00 95% [1.00, 1.06]), donor (**Figure 5B,E**, median iLISI 1.42 95% [1.00, 5.50]), and cell type (**Figure 5E**, median cLISI 1.00 95% [1.00, 1.48]). The wide range of donor-iLISI reflects that in the CEL-seq, CEL-seq2, and Fluidigm C1 datasets, many donors were well mixed prior to integration. Harmony integrates cells by both technology (**Figure 5C,E**, median iLISI 2.27 95% [1.31, 3.27]) and donor (**Figure 5D,E**, median iLISI 4.71 95% [1.81, 6.36]).

**Figure 5:**
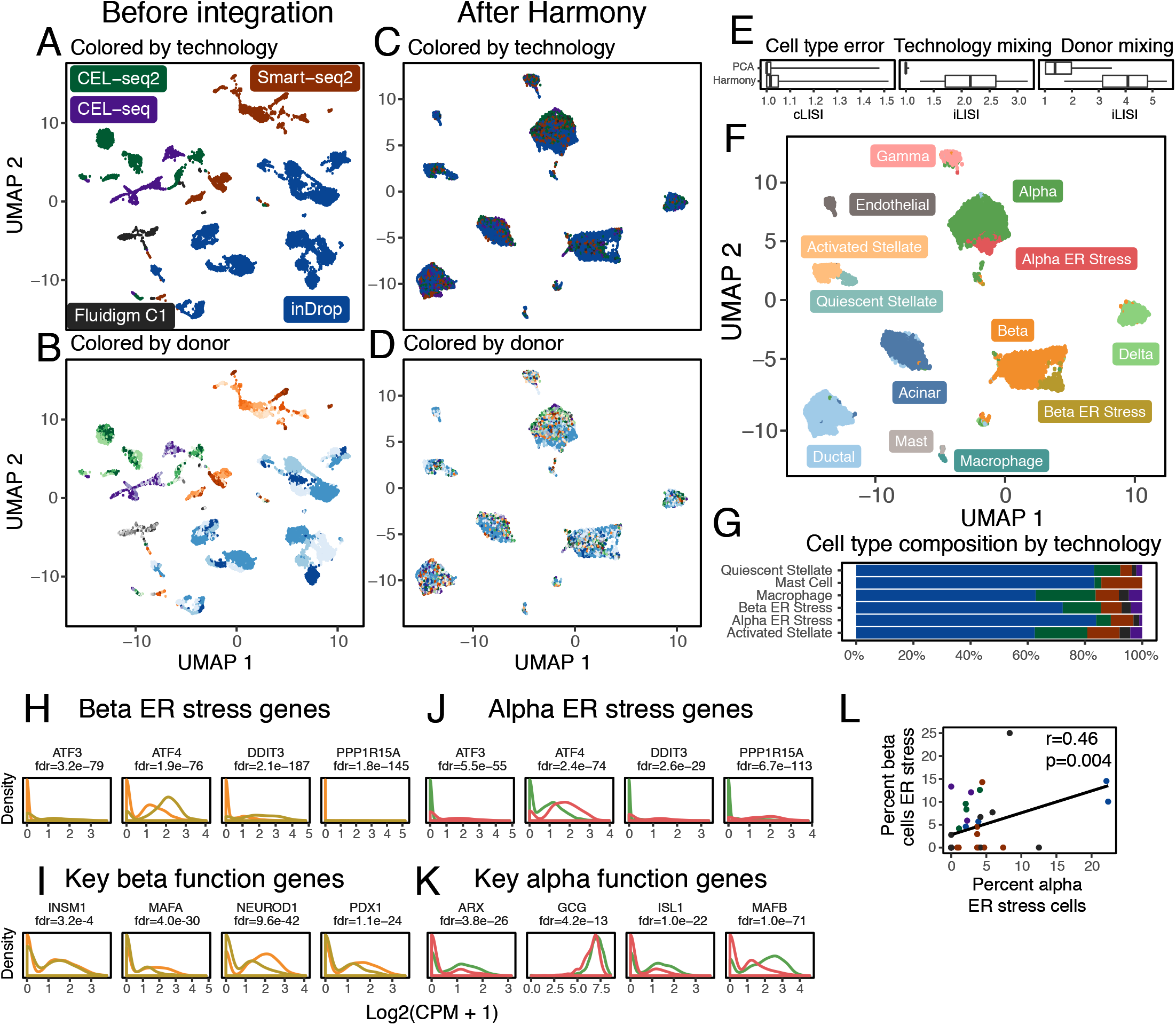
Integration of pancreatic islet cells by both donor and technology. Human pancreatic islet cells from 36 donors were assayed on 5 different technologies. Cells initially group by (A) technology, denoted by different colors, and (B) donor, denoted by shades of colors. Harmony integrates cells simultaneously across (C) technology and (D) donor. Integration across both variables was quantified with iLISI (E) and error was computed with cLISI (E). Clustering in the Harmony embedding identified common and rare cell types, including a previously undescribed alpha population. Except for activated stellate cells, all rare cell types were found across the 5 technology datasets (G). The new alpha cluster was enriched for ER stress genes (I), just like the previously identified beta ER stress cluster (J). The abundances of the two ER stress populations were correlated across donors (H). Key genes necessary for endocrine function were downregulated in the alpha (K) and beta (L) ES stress clusters.

Harmony was able to discern rare cell subtypes (**Figure 5F**) across the 5 datasets (**Figure 5G**). We labeled previously described subtypes using canonical markers: activated stellate cells (*PDGFRA*+), quiescent stellate cells (*RGS5*+), mast cells (*BTK*+), macrophages (*C1QC*+), and beta cells under endoplasmic reticulum (ER) stress (**Figure 5H**). Beta ER stress cells may represent a dysfunctional population. This cluster has significantly lower expression of genes key to beta cell identify^17^ and function:^18^ *PDX1*, *MAFA*, *INSM1*, *NEUROD1* (**Figure 5I**). Further, Sachdeva et al^19^ suggest that *PDX1* deficiency makes beta cells less functional and exposes them to ER stress induced apoptosis.

Intriguingly, we also observed an alpha cell subset that to our knowledge, has not been previously described. This cluster was also enriched with genes involved in ER stress (**Figure 5J**, *DDIT3*, *ATF3*, *ATF4*, and *HSPA5*). Similar to the beta ER stress population, these alpha cells also expressed significantly lower levels of genes necessary for proper function:^20, 21^ *GCG*, *ISL1*, *ARX*, and *MAFB* (**Figure 5K**). A recent study^22^ reported ER stress in alpha cells in mice and linked the stress to dysfunctional glucagon secretion. Moreover, we found that the proportions of alpha and beta ER stress cells are significantly correlated (spearman r=0.46, p=0.004, **Figure 5L**) across donors in all datasets. These results suggest a basis for alpha cell injury that might parallel beta cell dysfunction in humans during diabetes.^23^

## Discussion

We showed that Harmony address the three key challenges we laid out for single cell integration analyses: scaling to large datasets, identification of both broad populations and fine-grained subpopulations, and flexibility to accommodate complex experimental design. We evaluated the degree of mixing among datasets using a quantitative metric, the iLISI. Apart from its use benchmarking, iLISI was particularly important in analyses with more than 3 datasets. Here, we observed that the commonly utilized approach of assessing integration visually was subjective and insensitive, particularly when the number of samples, batches or cell types was large. iLISI provides a quantitative and interpretable metric to help guide analysis. In the computational efficiency benchmarks, we found that 3 of the 5 algorithms were not able to scale beyond 125,000 cells because they exceeded the memory resources of our 128GB servers. We were struck by the fact many researchers routinely analyze data on personal computers, which often do not exceed 8 or 16GB. Harmony, which only required 7.2GB to integrate 500,000 cells, is the only algorithm that would enable the integration of large datasets on personal computers. With the pancreatic islet meta-analysis, we demonstrated that Harmony is able to account simultaneously for donor and technology specific effects. One solution to this multi-level problem stepwise is to first globally regress out one variable from the gene expression values and then performing single cell integration on the resulting expression matrix. Harmony allows for cell-type aware integration of both variables, simultaneously avoiding global correction terms that treat all cells uniformly. However, the global regression strategy is flexible enough to account for continuous variables, such as read depth or cell quality. In the future, Harmony should also be able to account for such non-discrete sources of variation.

We noticed that it is not an uncommon practice to apply a batch-sensitive gene scaling step step before using a single-cell integration algorithm. Specifically, many investigators scale gene expression values within datasets separately, before pooling cells into a single matrix. We show (Supplementary Results) that this strategy may make it easier to integrate datasets (**Figure S8A,B**) in the rare situation in which all cell populations and subpopulations are present across all analyzed datasets. However, when the datasets consist of overlapping but not identical populations, this scaling strategy is less effective (**Figure S8C**) and may indeed even increase error (Figure S8D). For this reason, we do not use this scaling strategy in this manuscript. As part of a universal pipeline, Harmony finds highly integrated embeddings without the need for within-dataset scaling.

Harmony allows the user to set a hyperparameter for each covariate that guides how deeply to integrate over each source of variation. When the penalty is 0, Harmony performs minimal integration. Curiously, we noticed that for small inter-datasets differences, cells from multiple datasets cluster together without the penalty. In this case, Harmony still integrates the cells during the linear correction phase. On the other hand, one could imagine that with an infinitely large penalty hyperparameter, Harmony would overmix datasets during clustering and hence overcorrect the data. We evaluated the effect of the diversity penalty in the PBMCs example (see Supplementary Results) and observed that the Harmony embedding is robust to a wide range of penalties (**Figure S9**). Nonetheless, as with any integration algorithm, we urge the user to understand the effects of hyperparameters and experiment with several values.

Harmony is designed to accept a matrix of cells and covariate labels for each cell as input, and it will output a matrix of adjusted coordinates with the same dimensions as the input matrix. As such, Harmony should be used as an upstream step in a full analysis pipeline. Downstream analyses, such as clustering, trajectory analysis, and visualization, can use the integrated Harmony adjusted coordinates instead of the commonly used PCA coordinates. As a corollary, Harmony does not alter the expression values of individual genes to account for dataset-specific differences. We recommend using a batch-aware approach, such as a linear model with covariates, for differential expression analysis.

In our meta-analysis of pancreatic islet cells, we identified a previously undescribed rare subpopulation of alpha ER stress cells (**Figure 5F,J**). Similar to beta ER stress cells, they appear to have reduced endocrine function (**Figure 5K**). Because Harmony integrated over both donors and technology (**Figure 5C,D,E**), we were able to identify the significant association between the proportion of alpha to beta ER stress populations across donors (**Figure 5L**). Based on this correlation and similar stress response patterns, it is possible that these two populations are involved in a coordinated response to an environmental stress. Beta cell dysfunction is key to the pathogenesis of diabetes23. Experimental follow up on this alpha subtype and its relation to beta ER stress cells may yield insight into disease. This analysis demonstrates the power of Harmony’s multilevel integration to mix diverse datasets and uncover potentially novel rare cell types.

## Online Methods

### Harmony

The Harmony algorithm inputs some embedding (*Z*) of cells, along with their batch assignments (*ϕ*), and returns a batch corrected embedding (*Ẑ*). This algorithm, summarized as **Algorithm 1** below, iterates between two complementary stages: maximum diversity clustering (**Algorithm 2**) and a mixture model based linear batch correction (**Algorithm 3**). The clustering step uses the a batch corrected embedding *Ẑ* to compute a soft assignment of cells to clusters, encoded in the matrix *R*. The correction step uses these soft clusters to compute a new corrected embedding from the original one. Efficient implementations of Harmony, including the clustering and correction subroutines, are available as part of an R package at https://github.com/immunogenomics/harmony.

#### Algorithm 1 Harmony

~~~
      **function** HARMONIZE(*Z, ϕ*^(1)^ *… ϕ* ^(*F*)^)
         *Ẑ* ← *Z*
         **repeat**
            *R*← CLUSTER(*Ẑ, ϕ*^(1)^ *… ϕ*^(*F*)^)
            *Ẑ* CORRECT(*Z, R, ϕ*^(1)^)
         **until** convergence
         **return** *Ẑ*
~~~

Note that the correction procedure uses *Z*, not *Ẑ* to regress out confounder effects. In this way, we restrict correction to a linear model of the original embedding. An alternative approach would use the output *Ẑ* of the last iteration as input to the correction procedure. Thus, the final *Ẑ* would be the result of a series of linear corrections of the original embedding. While this allows for more expressive transformations, we found that in practice, this can over correct the data. Our choice to limit the transformation reflects the notion in the introduction. Namely, if we had perfect knowledge of the cell types before correction, we would linearly regress out batch within each cell type.

Lastly, we note that *Z* can be any arbitrary embedding of the cells. In this paper, we use principal components for scRNAseq datasets. However, Harmony can be efficiently run on any low dimensional embedding of the data, including diffusion maps, autoencoders, or independent components.

#### Glossary

For reference, we define all data structures used in all Harmony functions. For each one, we define its dimensions and possible values, as well as an intuitive description of what it means in context. The dimensions are stated in terms of **d**: the dimensionality of the embedding (e.g. number of PCs), **B**: the number of batches, **N**: the number of samples, **N_b_**: the number of samples in batch *b*, and **K**: the number of clusters.

*Z* ∈ ℝ^*d×N*^. The input embedding, to be corrected in Harmony. This is often PCA embeddings of cells.

*Ẑ* ∈ ℝ^*d×N*^. The integrated embedding, output by Harmony.

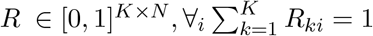. The soft cluster assignment matrix of cells (columns) to clusters (rows). Each column is a probability distribution and thus sums to 1.

*ϕ* ∈ {0, 1}^*B×N*^ One-hot assignment matrix of cells (columns) to batches (rows).

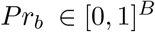. Frequency of batches.

*O* ∈ [0, 1]^*K×B*^ The observed co-occurence matrix of cells in clusters (rows) and batches (columns).

*E* ∈ [0, 1]^*K×B*^ The expected co-occurence matrix of cells in clusters and batches, under the assumption of independence between cluster and batch assignment.

*Y* ∈ [0, 1]^*d×K*^ Cluster centroid locations in the kmeans clustering algorithm.

*μ* ∈ ℝ^*d×K*^ Centroid locations. Conceptually the same as *Y* but not normalized to unit length. Used in GMM Correct.

*β* ∈ ℝ^*d×K×B*^ Batch-specific centroid locations. Used in GMM Correct.

Note that in the case of multiple (*F*) batch variables, *ϕ*, *θ*, *O*, and *E* are independently defined for each batch *f*. We denote these by *ϕ*^(*f*)^, *θ_f_*, *O*^(*f*)^, and *E*^(*f*)^ for each batch from 1 to *F*.

### Maximum Diversity Clustering

We developed a clustering algorithm to maximize the diversity among batches within clusters. We present this method as follows. First, review a previously published objective function for soft k-means clustering.

We then add a diversity maximizing regularization term to this objective function, and derive this regularization term as the penalty on statistical dependence between two random variables: batch membership and cluster assignment. We then derive and present pseudocode for an algorithm to optimize the objective function. Finally, we explain key details of the implementation.

#### Background: Entropy regularization for Soft K-means

The basic objective function for classical K means clustering, in which each cell belongs to exactly one cluster, is defined by the distance from cells to their assigned centroids.

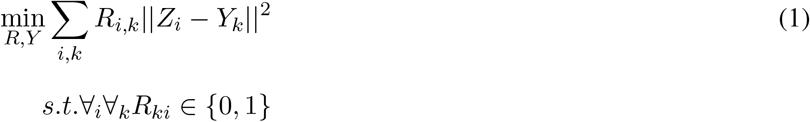

Above, *Z* is some feature space of the data, shared by centroids *Y*. *R_i,k_* can takes values 0 or 1, denoting membership of cell *i* in cluster *k*. In order to transform this into a soft clustering objective, we follow the direction of^24^ and add a an entropy regularization term over *R*, weighted by a hyperparameter *σ*. Now, *R_ki_* can take values between 0 and 1, so long as for a given cell *i*, the sum over cluster memberships Σ*_k_ R_ki_* equals 1. That is, *R_i_*. must be a proper probability distribution with support [1*, K*].

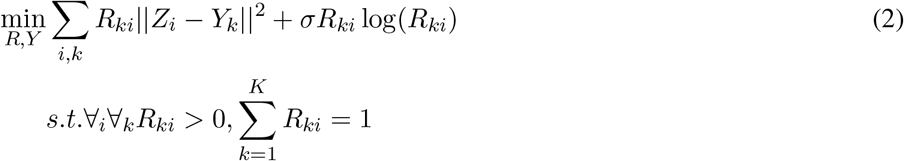

As *σ* approaches 0, this penalty approach hard clustering. As *σ* approaches infinity, the entropy of *R* outweighs the data-centroid distances. In this case, each data point is assigned equally to all clusters.

#### Objective Function for Maximum Diversity Clustering

The full objective function for Harmony’s clustering builds on the previous section. In addition to soft assignment regularization, the function below penalizes clusters with low batch-diversity, for all defined batch variables. This penalty, derived in the following section, depends on the cluster and batch identities 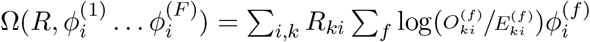.

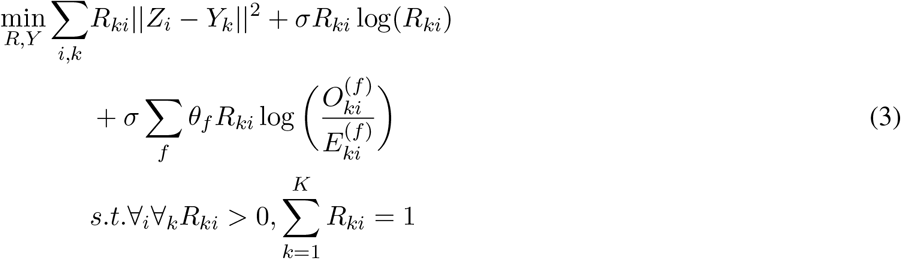

For each batch variable, we add a new parameter *θ_f_*. *θ_f_* decides the degree of penalty for dependence between batch membership and cluster assignment. When ∀*_f_ θ_f_* = 0, the problem reverts back to (2), with no penalty on dependence. As *θ_f_* increases, the objective function favors more independence between batch *f* and cluster assignment. As *θ_f_* approaches infinity, it will yield a degenerate solution. In this case, each cluster has an equivalent distribution across batch *f*. However, the distances between cells and centroids may be large. Finally, *σ* is added to this term for notational convenience in the gradient calculations.

We found that this clustering works best when we compute the cosine, rather than Euclidean distance, between *Z* and *Y*. Haghverdi et al^25^ showed that the squared Euclidean distance is equivalent to cosine distance when the vectors are *L*_2_ normalized. Therefore, assuming that all *Z_i_* and *Y_k_* have a unity *L*_2_ norm, the squared Euclidean distance above can be re-written as a dot product.

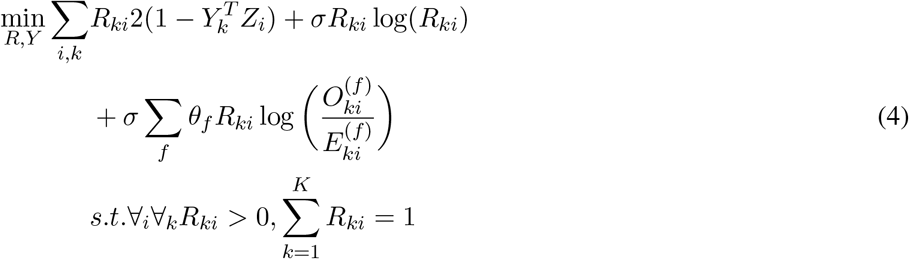

#### Cluster Diversity Score

Here, we discuss and derive the diversity penalty term Ω(⋅), defined in the previous section. For simplicity, we discuss diversity with respect to a single batch variable, as the multiple batch penalty terms are additive in the objective function. The goal of Ω(⋅) is to penalize statistical dependence between batch identity and cluster assignment. In statistics, dependence between two discrete random variables is typically measured with the *χ*^2^ statistic. This test considers the frequencies with which different values of the two random variables are observed together. The observed co-occurrence counts (*O*) are compared to the counts expected under independence (*E*). For practical reasons, we do not use the *χ*^2^ statistic directly.

Instead, we use the Kullback Leibler Divergence (*D_KL_*), an information theoretic distance between two distributions. In this section, we define the *O* and *E* distributions, as well the *D_KL_* penalty, in the context of the probabilistic cluster assignment matrix *R*.

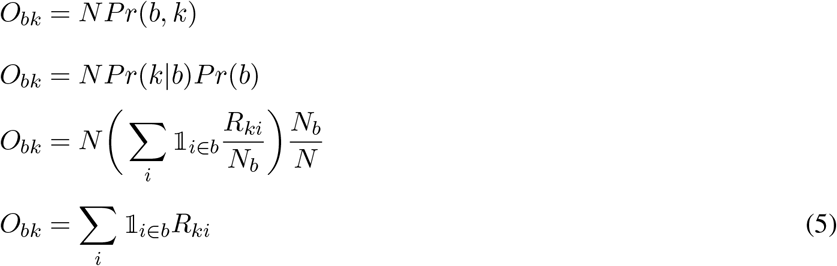

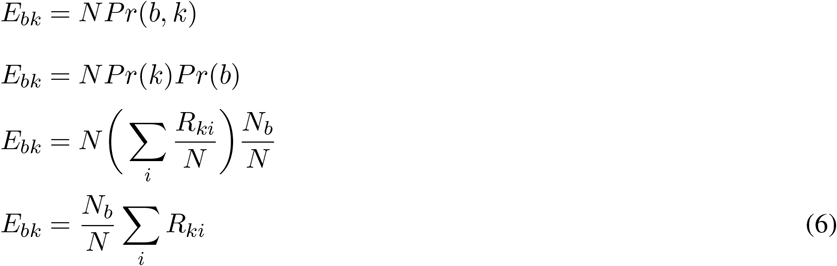

Next, we define the KL divergence in terms of *R*. Note that both *O* and *E* depend on *R*. However, in the derivation below, we expand one of the *O* terms. This serves a functional purpose in the optimization procedure, described later. Intuitively, in the update step of *R* for a single cell, we compute *O* and *E* on all the other cells. In this way, we decide how to assign the single cell to clusters given the current distribution of batches amongst clusters.

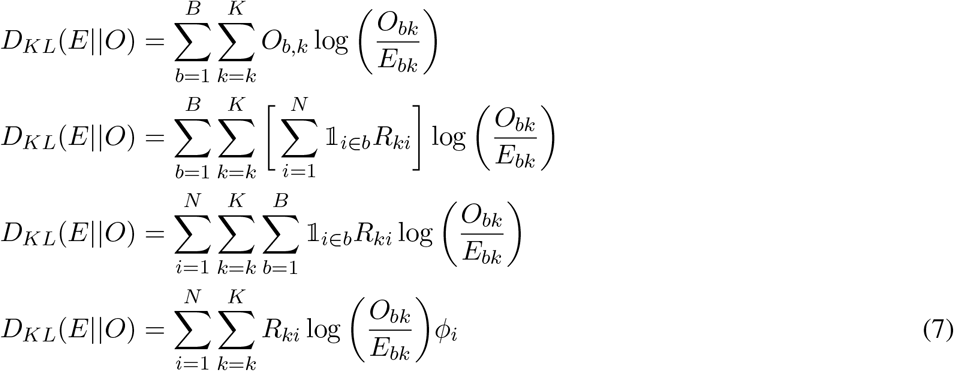

For multiple batch variables, we sum over these convergence terms to get the penalty in equation 3.

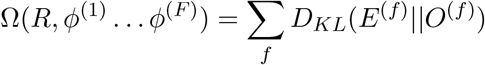

#### Optimization

Optimization of 4 admits an Expectation-Maximization framework, iterating between cluster assignment (*R*) and centroid (*Y*) estimation.

##### Cluster assignment *R*

Using the same strategy as,^24^ we solve for the optimal assignment *R_i_* for each cell *i*. First we set up the Lagrangian with dual parameter *λ* and solve for the partial derivative wrt each cluster *k*.

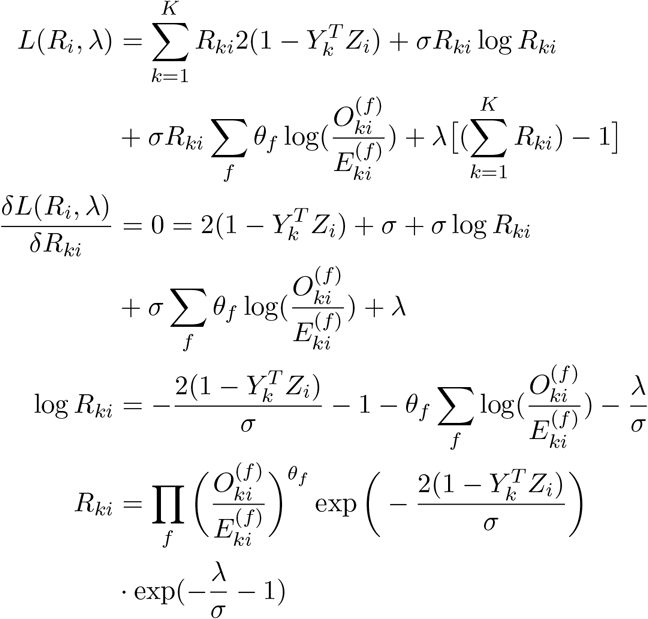

Next, we use the probability constraint 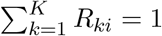 to solve for 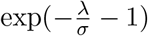.

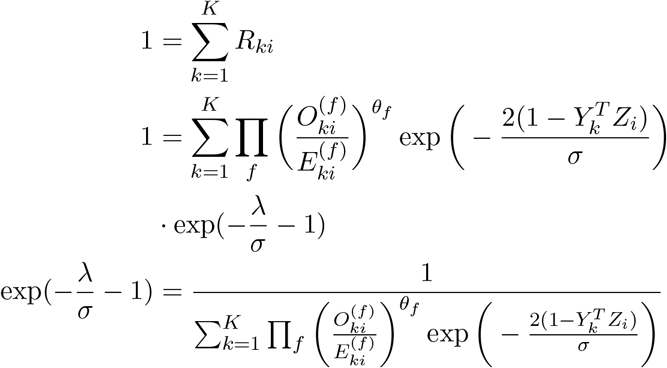

Finally, we substitute 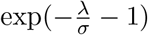 to remove the dependency of *R_ki_* on the dual parameter *λ*.

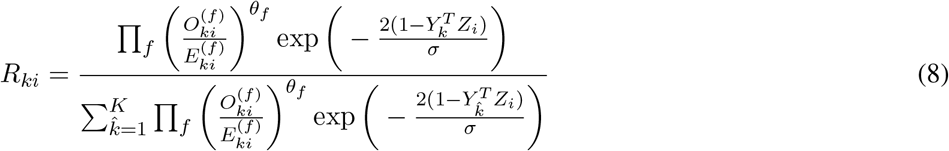

The denominator term above makes sure that *R_i_* sums to one. In practice (alg 2), we compute the numerator and divide by the sum.

##### Centroid Estimation *Y*

Our clustering algorithm uses cosine distance instead of Euclidean distance. In the context of kmeans clustering, this approach was pioneered by Dhillon et al.^26^ We adopt their centroid estimation procedure for our algorithm. Instead of just computing the mean position of all cells that belong in cluster *k*, this approach then *L*_2_ normalizes each centroid vector to make it unit length. Note that normalizing the sum over cells is equivalent to normalizing the mean of the cells. In the soft clustering case, this summation is an expected value of the cell positions, under the distribution defined by *R*. That is, re-normalizing *R_.k_* for cluster *k* gives the probability of each cell belonging to cluster *k*. Again, this re-normalization is a scalar factor that is irrelevant once we *L*_2_ normalize the centroids. Thus, the unnormalized expectation of centroid position for cluster *k* would be *Y_k_* = 𝔼*_R_._k_ Z* = Σ*_i_ R_ki_Z_i_*. In vector form, for all centroids, this is *Y* = *ZR^T^*. The final position of the cluster centroids is given by this summation followed by *L*_2_ normalization of each centroid. This procedure is implemented in algorithm 2 in the section {*Compute Cluster Centroids*}.

###### Algorithm 2 Maximum Diversity Clustering

~~~
     **function** CLUSTER(*Ẑ, ϕ*^(1)^ *… ϕ*^(*F*)^)
          {*Initialize Cluster Centroids*}
          *Y ←* **0**_[*d×K*]_
          **for** *b ←* 1*…B* **do**
                 *sample ←* K random points from batch b
                 *Y ← Y* + *Z_sample_*
          **for** *i ←* 1*…N* **do**                  *⊳ L*_2_ Normalization
                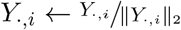
               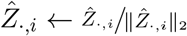
          **for** f←1…*F* **do**
               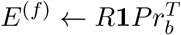
               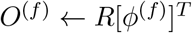
          **repeat**
              **for all** Update Blocks **do**
               *in ←* cells to update in block
               {*Compute O and E on left out data*}
               **for** *f ←* 1*…F* **do**
                     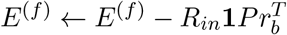
                     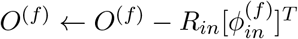
               {*Update and Normalize R*}
               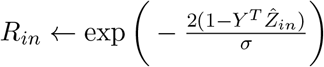
               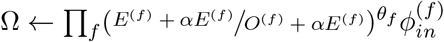
               *R_in_ ← R_in_ ∘* Ω
               *R_in_ ← Rin/***1***^T^ R_in_                   ⊲ R_i_* sum to one
               {*Compute O and E with full data*}
               **for** *f ←* 1*…F* **do**
                    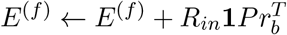
                    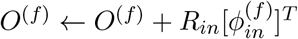
            {*Compute Cluster Centroids*}
            *Y ← ZR^T^*
            **for** *i ←* 1*…N* **do**                      *⊲L*_2_ Normalization
                    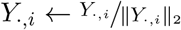
        **until** convergence
        **return** *R*
~~~

### Implementation Details

The update steps of *R* and *Y* derived above form the core of Maximum Diversity Clustering, outlined as algorithm 2. This section explains the other implementation details of this pseudocode. Again, for simplicity, we discuss details related to diversity penalty terms *θ*, *φ*, *O*, and *E* for each single batch variable independently.

#### Block Updates of *R*

Unlike in regular kmeans, the optimization procedure above for *R* cannot be faithfully parallelized, as the values of *O* and *E* change with *R*. The exact solution therefore depends an online procedure. For speed, we can coarse grain this procedure and update *R* in small blocks (e.g. 5% of the data). Meanwhile, *O* and *E* are computed on the held out data. In practice, this approach succeeds in minimizing the objective for sufficiently small block size. In the algorithm, these blocks are included as the Update Blocks in the for loop.

#### Centroid Initialization

We initialize cluster centroids using a batch balanced random medeoid approach. We randomly select *K* cells within each batch for a total of *B × K* centroids. We then randomly select one centroid from each batch, average it, and *L*_2_ normalize it to create a total of *K* centroids. With this strategy, any single centroid is unlikely to be centered within one batch.

### Regularization for Smoother Penalty

The diversity penalty term (*E_bk_/O_bk_*)^*θ*^ can tend towards infinity if there are no cells from batch *b* assigned to cluster *k*. This extreme penalty can erroneously force cells into an inappropriate cluster. To protect against this, we smooth this term to ensure that the denominator will not be zero. The strategy is to use *E* as a prior and add some portion *α* ̂ [0, 1] of it to the empirically estimated values of *E* and *O*. As a result, the smoothed fraction becomes 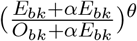.

#### *θ* Discounting

The diversity penalty, weighted by *θ* enforces an even mixing of cells from a batch among all clusters. This assumption is more likely to break for a batch with few cells. The smaller the batch, the more likely it is, through a sampling argument, that some cell types are not represented in the batch. Spreading such a batch across all clusters would result in erroneous clustering. To prevent such a situation, we allot each batch its own *θ_b_* term, scaled to the number of cells in the batch.

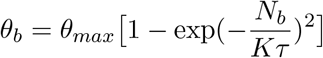

Above, *θ_max_* is the non-discounted *θ* value, for a large enough batch. The multiplicative factor 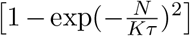 ranges from 0 to 1. This factor scales exponentially for small values of batch size *N_b_* and plateaus for sufficiently large *N_b_*. The hyperparameter *τ* can be interpreted as the minimum number of cells that should be assigned to each cluster from a single batch. By default, we use values between *τ* = 5 and *τ* = 20.

### Linear Mixture Model Correction

In this section, we refer to all effects to be integrated out of the original embedding as batch effects. This does not imply that these effects are purely technical. This terminology is only meant for convenience. After clustering, we estimate and correct for additive batch effect in low dimensional space. The intuition behind our approach is to give each cell within a batch a different batch effect term, depending on its cell type/state. As we do not know the latter a priori, we use cluster membership as a surrogate variable. For simplicity, we choose to limit our treatment of batch effect to the additive mode and avoid estimating scaling effects. Note that although there are potentially multiple batch assignments handled in the clustering above, the regression step currently deals with only one variable. Here, we assume that *ϕ* is *ϕ*^(1)^.

#### Simple Additive Batch Model

. Before we jump in to the full mixture model, we review a simpler model of additive batch. Here, batch effect is global and is not sensitive to any surrogate variables, such as cluster membership. We assume the data is normally distributed and model each sample (*i*) with a multivariate Gaussian.

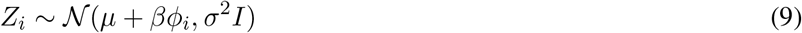

The covariance matrix Σ is a diagonal matrix with the same variance *σ*^2^ for each dimension. The mean is a sum of a global intercept vector *μ* and a batch effect vector *βϕ_i_*. In this example, *β* ∈ ℝ^*d×B*^ defines an offset vector for each batch. We use *φ_i_* to index into the appropriate batch of *β* (i.e. *βϕ_i_* ∈ ℝ^*d×*1^). In order to regress out the batch effect, we estimate the parameters *μ* and *β* and subtract the batch offsets (*βϕ_i_*) from the raw data. Since batch effect here is purely additive, we do not estimate variance. *μ* is the global mean of the dataset 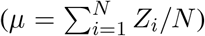. *β* terms are estimated by maximizing the log likelihood of (9) wrt each *β_b_* term independently.

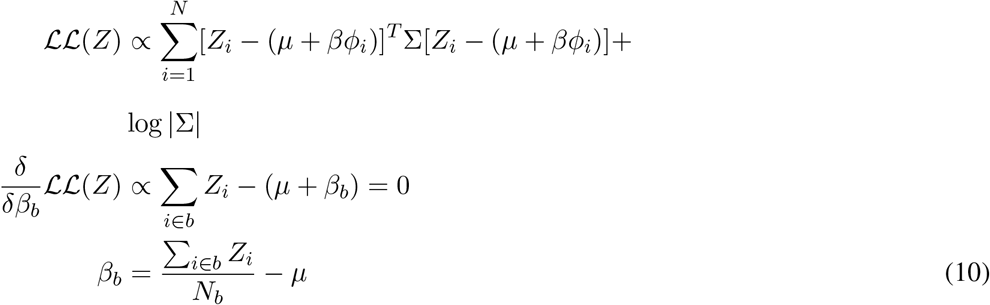

Thus, the batch offset of batch *b* is the mean of cells inside the batch minus the global mean. We subtract this offset from the raw data (*Z*) to get the corrected data (*Ẑ*).

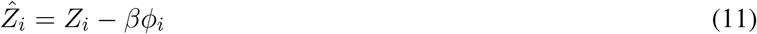

In the following section, we follow the same approach laid out here to incorporate soft cluster membership into the model.

#### Additive Batch Mixture Model

In order to model the effect of batch, we take advantage of the parameterization of K-means as a special case of a Gaussian mixture model (GMM). That is, the cluster assignment probabilities (*R*) computed before now become the component mixture weights in the GMM. Like above, this GMM has a homoschedastic, spherical covariance matrix (Σ = *σ*^2^*I*). Unlike the simpler model, each cluster has its own intercept *μ_k_* and set of batch terms *β_bk_*.

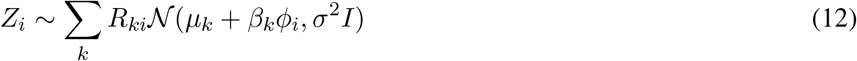

In this new generative model, each sample is explained as a mixture of Gaussians, with weights *R_ki_*. The cluster specific intercepts (*μ_k_*) are the (non-spherical) cluster centroids. We compute the MLE *β* terms using standard methods for GMMs. It is known that the GMM does not have a convex log likelihood, so we use the the expected log likelihood instead. Fixing *μ_k_* and *R_ki_*, we solve for the MLE *β_bk_* terms.

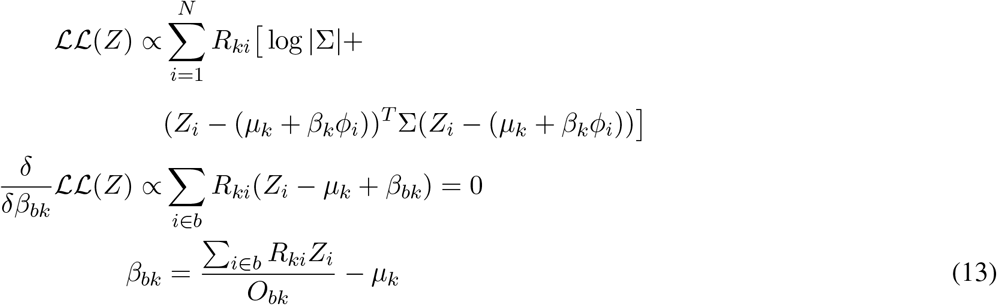

Note that (10) and (13) have similar forms. Both compute the expected difference between the batch cells and some *μ*. In the simple model, *μ* is a global term, while in the mixture model, *μ_k_* is specific to a local cluster *k*. In order to correct the data in this model, we need a batch offset term for each cell. (12) shows us that this is done by taking expectations over *β_k_ϕ_i_* terms:

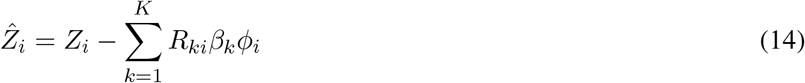

#### Implementation

Algorithm 3 lays down the pseudocode for GMM Correct, the mixture model correction procedure derived above. For a more vectorized implementation, we rewrite the (13) in terms of each *β_b_* ∈ ℝ^*d×K*^ matrix and (14) in terms of the whole *Z* matrix.

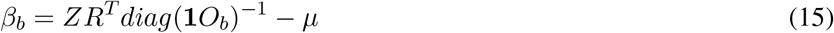

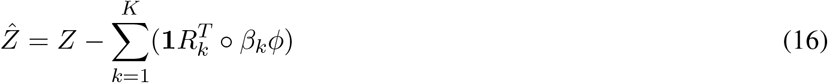

##### Algorithm 3 GMM Correct

~~~
    **function** CORRECT(*Z, R, ϕ*)
          {*Compute Cluster Means*}
          Γ *← diag*(1*^T^ R*)
          *μ ← ZR^T^* Γ^−1^*R*
          {*Compute Batch Offsets*}
          *O_bk_ ← Rϕ^T^*
          *β ←* **0**_[*d×K×B*]_
          **for** *b ←* 1 *… B* **do**
                Γ*_b_ ← diag*(1*O_b_*)
               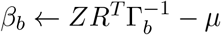
          {*Compute Batch Corrected Output}*
          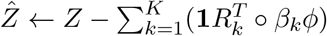
          **return** *Ẑ*
~~~

#### Caveat

This section assumes the modeled data are orthogonal and each normally distributed. This is not true for the *L*_2_ normalized data used in spherical clustering. Regression in this space requires the estimation and interpolation of rotation matrices, a difficult problem. We instead perform batch correction in the unnormalized space. The corrected data *Ẑ* are then *L*_2_-normalized for the next iteration of clustering.

### Performance and Benchmarking

#### LISI Metric

Assessing the degree of mixing during batch correction and dataset integration is an open problem. Several groups have proposed methods to quantify the diversity of batches within local neighborhoods, defined by k nearest neighbor (KNN) graphs, of the embedded space. Buttner et al^27^ provide a statistical test to evaluate the degree of mixing, while Azizi et al^28^ report the entropy of these distributions. Our metric for local diversity is related to these approaches, in that we start with a KNN graph. However, our approach considers two problems that these do not.

First, the metric should be more sensitive to local distances. For example, a neighborhood of 100 cells can be equally mixed among 4 batches. However, within the neighborhood, the cells may be clustered by batch. The second problem is one of interpretation. kBET provides a statistical test to assess the significance of mixing, but it is not clear whether all neighborhood should be significantly mixed when the datasets have vastly different cell type proportions. Azizi et al^28^ et al use entropy as a measure of diversity, but it is not clear how to interpret the number of bits required to encode a neighborhood distribution.

Our diversity score, the Local Inverse Simpson’s Index (LISI) addresses both points. To be sensitive to local diversity, we build Gaussian kernel based distributions of neighborhoods. This gives distance-based weights to cells in the neighborhood and gives less diversity to. The current implementation computes these local distributions using a fixed perplexity (default 30), which has been shown to be a smoother function than fixing the number of neighbors. We address the second issue of interpretation using the Inverse Simpson’s Index 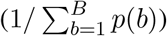. The probabilities here refer to the batch probabilities in the local neighborhood distributions described above. This index is the expected number of cells needed to sampled before two are drawn from the same batch. If the neighborhood consists of only one batch, then only one draw is needed. If it is an equal mix of two batches, two draws are required on average. Thus, this index gives the effective number of batches in a local neighborhood. Our diversity score, LISI, combines these two features: perplexity based neighborhood construction and the Inverse Simpson’s Index. LISI assigns a diversity score to each cell. This score is the effective number of batches in that cell’s neighborhood. Code to compute LISI is available as at https://github.com/immunogenomics/LISI.

#### Time and Memory

We performed execution time and maximum memory usage benchmarks on all analyses. All jobs were run on Linux servers and allotted 6 cores and 120GB of memory. The machines were equipped with Intel Xeon E5-2690 v3 processors. To evaluate execution time and maximal memory usage, we used the Linux time utility (/usr/bin/time on our systems) with the -v flag to record memory usage. Execution time was recorded from the *Elapsed time* field. Maximum memory usage was recorded from the *Maximum resident set size* field.

### Analysis Details

#### Data Availability

All data analyzed in this manuscript is publicly available through online sources. We included links to all data sources in **Table S6.**

#### Preprocessing scRNAseq data

We downloaded raw read or UMI matrices for all datasets, from their respective sources. The one exception was the 3pV1 dataset from the PBMC analysis. These data were originally quantified with the hg19 reference, while the other two PBMC datasets were quantified with GRCh38. Thus, we downloaded the fastq files from the 10X website (**Table S6**). We quantified gene expression counts using Cell Ranger^10, 29^ v2.1.0 with GRCh38. From the raw count matrices, we used a standard data normalization procedure, laid out below, for all analyses, unless otherwise specified. Except for the *L*_2_ normalization and within-batch variable gene detection, this procedure follows the standard guidelines of the Seurat single cell analysis platform.

We filtered cells with fewer than 500 genes or more than 20% mitochondrial reads. In the pancreas datasets, we filtered cells with the same thresholds used in Butler et al:6 1750 genes for CelSeq, 2500 genes for CelSeq2, no filter for Fluidigm C1, 2500 genes for SmartSeq2, and 500 genes for inDrop. We then library normalizes each cell to 10,000 reads, by multiplicative scaling, then log scaled the normalized data. We then identified the top 1000 variable genes, ranked by coefficient of variation, within in each dataset. We pooled these genes to form the variable gene set of the analysis. Using only the variable genes, we mean centered and variance 1 scaled the genes across the cells. Note that this was done in the aggregate matrix, with all cells, rather than within each dataset separately. We then *L*_2_ normalized the cell expression vectors, so that squared cosine distance can be computed as the squared Euclidean distance. With these values, performed truncated SVD keeping the top 30 eigenvectors. Finally, we multiplied the cell embeddings by the eigenvalues to avoid giving eigenvectors equal variance.

#### Visualization

We used the UMAP algorithm^30^ to visualize cells in a two dimensional space. For all analyses, UMAP was run with the following parameters: *k* = 30 nearest neighbors, correlation based distance, and *min dist* = 0.1.

#### Comparison to other algorithms

We used the provided packages or source code provided by the four comparison algorithm publications. For MNN Correct, we used the mnncorrect function, with default parameters, in the scran R package,^31^ version 1.9.4. For Seurat MultiCCA, we followed the suggested integration pipeline in the Seurat R package,^32^ version 2.3.4. Specifically, this included the RunMultiCCA, MetageneBicorPlot, CalcVarExpRatio, SubsetData (based on the var.ratio.pca statistic), and AlignSubspace functions. We used all default parameters. We chose the number of canonical components to match the number of principal components used in the preprocessing. Unlike the integration examples, we did not scale data within datasets separately, unless otherwise specified. We scaled data on the pooled count matrix instead. For Scanorama, we used the assemble function, with precomputed PCs, from the primary github repository (brianhie/scanorama). We set knn=30 and sigma=1, to match the default comparable MNN Correct parameters. All other parameters were kept at default values. We did not use the correct function, as this included both pre-processing and integration of the data. For more equitable comparisons, we tried to use the same pre-processing pipelines for all methods and only compare only the integration steps. For BBKNN, we downloaded software from the primary github repository (Teichlab/bbknn) and followed the suggested integration pipelines, using the bbknn and scanpy umap functions. For the bbknn function, we used k=5 and trim=20 for all analyses except for the HCA datasets, in which we used k=10 and trim=30, to accommodate the larger number of cells. All other parameters were kept at default values.

#### Harmony Parameters

By default, we set the following parameters for Harmony: *θ* = 2, *K* = 100, *τ* = 0, *α* = 0.1, *σ* = 0.1 block_size= 0.05, *E_cluster_* = 10^*−*5^, *E_harmony_* = 10^*−*4^, max_iter_*cluster*_ = 200, max_iter_*Harmony*_ = 10. For the pancreas analysis, we set *τ* = 5. We set donors to be the primary covariate (*θ* = 2) and technology secondary (*θ* = 4).

#### Identification of alpha and beta ER stress subpopulations

We identified the alpha and beta ER stress clusters in **Figure 5** by performing downstream analysis, specified in this section, on the integrated joint embedding produced by Harmony. After Harmony integration, we performed clustering analysis to find novel subtypes. Clustering was done on the trimmed shared nearest neighbor graph with the Louvain algorithm,^33^ as implemented in the Seurat package BuildSNN and RunModularityClustering functions. We used parameters resolution=0.8, k=30, and nn.eps=0. We identified several clusters within the alpha, beta, and ductal cell populations. For each cluster, we performed differential expression analysis within the defined cell type. That is, we compared alpha clusters to all other alpha cells. For differential expression, we used the R Limma package^34^ on the normalized data. We included technology and library complexity (log number of unique genes) as covariates in the linear models. We used the top 100 over-expressed genes for each cluster, weighted by the t-statistic, to perform pathway enrichment with the enrichR^35, 36^ R package, using the three Gene Ontology genesets.^37, 38^ The ductal subpopulation was enriched for ribosomal genes; we did not follow up on this cluster. The results for the differential expression and enrichment analyses for the two ER stress subpopulations is available in Tables S7 through S10.

#### Labeling cells with canonical markers

In the cell-line, PBMC, and Pancreas analyses, we labeled cells within individual datasets using canonical markers. We did this by using the standard pre-processing pipeline for each dataset, clustering (Louvain, as above), and identifying clusters specific for the canonical markers for that analysis. We used a similar strategy to identify fine-grained subpopulations of PBMCs and in the HCA 500,000 cell dataset. In these case, we clustered in the joint embedding produced by Harmony, then looked for clusters that specifically expressed expected canonical markers.

## Acknowledgements

This work was supported in part by funding from the National Institutes of Health (UH2AR067677, U19AI111224, and 1R01AR063759 (to S.R.)) and T32 AR007530-31. We thank members of the Raychaudhuri and Brenner labs for comments and discussion.

## Author Contributions

SR and IK conceived the research. IK led computational work under the guidance of SR, assisted by PL, JF, and KS. All authors participated in interpretation and writing the manuscript.

## Footnotes

^1^1.5 = 1 / [(1565 / (1565 + 2859))2 + (2859/ (1565 + 2859))2], 1.8 = 1 / [(1799 / (1799 + 2859))2 + (3255 / (1799 + 3255))2]

## References

1 Svensson, V., Vento-Tormo, R. & Teichmann, S. A. Exponential scaling of single-cell RNA-seq in the past decade. Nat. Protoc. 13, 599–604 (2018).

2 Regev, A. et al. The human cell atlas. Elife 6 (2017).

3 Zhang, F. et al. Defining inflammatory cell states in rheumatoid arthritis joint synovial tissues by integrating single-cell transcriptomics and mass cytometry. bioRxiv (2018).

4 Arazi, A. et al. The immune cell landscape in kidneys of lupus nephritis patients (2018).

5 Hicks, S. C., Townes, F. W., Teng, M. & Irizarry, R. A. Missing data and technical variability in single-cell RNA-sequencing experiments. Biostatistics (2017).

6 Butler, A., Hoffman, P., Smibert, P., Papalexi, E. & Satija, R. Integrating single-cell transcriptomic data across different conditions, technologies, and species. Nat. Biotechnol. 36, 411–420 (2018).

7 Haghverdi, L., Lun, A. T. L., Morgan, M. D. & Marioni, J. C. Batch effects in single-cell RNA-sequencing data are corrected by matching mutual nearest neighbors. Nat. Biotechnol. 36, 421–427 (2018).

8 Hie, B. L., Bryson, B. & Berger, B. Panoramic stitching of heterogeneous single-cell transcriptomic data (2018).

9 Park, J.-E., Polanski, K., Meyer, K. & Teichmann, S. A. Fast batch alignment of single cell transcriptomes unifies multiple mouse cell atlases into an integrated landscape (2018).

10 Zheng, G. X. Y. et al. Massively parallel digital transcriptional profiling of single cells. Nat. Commun. 8, 14049 (2017).

11 Li, B. et al. HCA data portal - census of immune cells.

12 Segerstolpe, Å. et al. Single-Cell transcriptome profiling of human pancreatic islets in health and type 2 diabetes. Cell Metab. 24, 593–607 (2016).

13 Baron, M. et al. A Single-Cell transcriptomic map of the human and mouse pancreas reveals inter-and intra-cell population structure. Cell Syst 3, 346–360.e4 (2016).

14 Lawlor, N. et al. Single-cell transcriptomes identify human islet cell signatures and reveal cell-type-specific expression changes in type 2 diabetes. Genome Res. 27, 208–222 (2017).

15 Grün, D. et al. De novo prediction of stem cell identity using Single-Cell transcriptome data. Cell Stem Cell 19, 266–277 (2016).

16 Muraro, M. J. et al. A Single-Cell transcriptome atlas of the human pancreas. Cell Syst 3, 385–394.e3 (2016).

17 Gao, T. et al. Pdx1 maintains *β* cell identity and function by repressing an *α* cell program. Cell Metab. 19, 259–271 (2014).

18 Jia, S. et al. Insm1 cooperates with neurod1 and foxa2 to maintain mature pancreatic *β*-cell function. EMBO J. 34, 1417–1433 (2015).

19 Sachdeva, M. M. et al. Pdx1 (MODY4) regulates pancreatic beta cell susceptibility to ER stress. Proc. Natl. Acad. Sci. U. S. A. 106, 19090–19095 (2009).

20 Katoh, M. C. et al. MafB is critical for glucagon production and secretion in mouse pancreatic *α* cells in vivo. Mol. Cell. Biol. 38 (2018).

21 Liu, J. et al. Islet-1 regulates arx transcription during pancreatic islet *α*-Cell development. J. Biol. Chem. 286, 15352–15360 (2011).

22 Akiyama, M. et al. X-box binding protein 1 is essential for insulin regulation of pancreatic *α*-cell function. Diabetes 62, 2439–2449 (2013).

23 Burcelin, R., Knauf, C. & Cani, P. D. Pancreatic alpha-cell dysfunction in diabetes. Diabetes Metab. 34 Suppl 2, S49–55 (2008).

24 Mao, Q., Wang, L., Goodison, S. & Sun, Y. Dimensionality reduction via graph structure learning. In Proceedings of the 21th ACM SIGKDD International Conference on Knowledge Discovery and Data Mining, KDD’15, 765–774 (ACM, New York, NY, USA, 2015).

25 Haghverdi, L., Lun, A. T. L., Morgan, M. D. & Marioni, J. C. Batch effects in single-cell RNA-sequencing data are corrected by matching mutual nearest neighbors. Nat. Biotechnol. 36, 421–427 (2018).

26 Dhillon, I. S. & Modha, D. S. Concept decompositions for large sparse text data using clustering. Mach. Learn. 42, 143–175 (2001).

27 Buttner, M., Miao, Z., Wolf, A., Teichmann, S. A. & Theis, F. J. Assessment of batch-correction methods for scRNA-seq data with a new test metric (2017).

28 Azizi, E. et al. Single-Cell map of diverse immune phenotypes in the breast tumor microenvironment. Cell 174, 1293–1308.e36 (2018).

29 Dobin, A. et al. STAR: ultrafast universal RNA-seq aligner. Bioinformatics 29, 15–21 (2013).

30 McInnes, L. & Healy, J. UMAP: Uniform manifold approximation and projection for dimension reduction. arXiv (2018). 1802.03426.

31 Lun, A. T. L., McCarthy, D. J. & Marioni, J. C. A step-by-step workflow for low-level analysis of single-cell rna-seq data with bioconductor. F1000Res. 5, 2122 (2016).

32 Butler, A., Hoffman, P., Smibert, P., Papalexi, E. & Satija, R. Integrating single-cell transcriptomic data across different conditions, technologies, and species. Nat. Biotechnol. (2018). URL https://www.nature.com/articles/nbt.4096.

33 Blondel, V. D., Guillaume, J.-L., Lambiotte, R. & Lefebvre, E. Fast unfolding of communities in large networks. J. Stat. Mech: Theory Exp. 2008 (2008).

34 Ritchie, M. E. et al. limma powers differential expression analyses for RNA-sequencing and microarray studies. Nucleic Acids Research 43, e47 (2015).

35 Chen, E. Y. et al. Enrichr: interactive and collaborative HTML5 gene list enrichment analysis tool. BMC Bioinformatics 14, 128 (2013).

36 Kuleshov, M. V. et al. Enrichr: a comprehensive gene set enrichment analysis web server 2016 update. Nucleic Acids Res. 44, W90–7 (2016).

37 The Gene Ontology Consortium. Expansion of the gene ontology knowledgebase and resources. Nucleic Acids Res. 45, D331–D338 (2017).

38 Ashburner, M. et al. Gene ontology: tool for the unification of biology. the gene ontology consortium. Nat. Genet. 25, 25–29 (2000).

